# Pathogenicity of SARS-CoV-2 Omicron in Syrian hamsters and its neutralization with different Variants of Concern

**DOI:** 10.1101/2022.01.19.477013

**Authors:** Sreelekshmy Mohandas, Pragya D. Yadav, Gajanan Sapkal, Anita M. Shete, Gururaj Deshpande, Dimpal A. Nyayanit, Deepak Patil, Manoj Kadam, Abhimanyu Kumar, Chandrashekhar Mote, Rajlaxmi Jain

## Abstract

SARS-CoV-2 Omicron variant is rampantly spreading across the globe. Animal models are useful in understanding the disease characteristics as well as properties of emerging SARS-CoV-2 variants. We assessed the pathogenicity and immune response generated by BA.1 sub-lineage of SARS-CoV-2 Omicron variant with R346K mutation in 5 to 6-week old Syrian hamsters. Virus shedding, organ viral load, lung disease and immune response generated were sequentially assessed. The disease characteristics of Omicron were found to be similar to that of other SARS-CoV-2 variants of concerns in hamsters like high viral replication in the respiratory tract and interstitial pneumonia. The infected hamsters demonstrated lesser body weight gain in comparison to the uninfected control hamsters. Viral RNA could be detected in nasal washes and respiratory organs (nasal turbinate, trachea, bronchi and lungs) till 10 and 14 days respectively. The clearance of the virus was observed from nasal washes and lungs by day 7. Neutralizing antibody response against Omicron variant was detected from day 5 with rising antibody titers till 14 days. However, the cross-neutralization titre of the sera against other variants showed severe reduction ie., 7 fold reduction against Alpha and no titers against B.1, Beta and Delta. This preliminary data shows that Omicron variant infection can produce moderate to severe lung disease and the neutralizing antibodies produced in response to Omicron variant infection shows poor neutralizing ability against other co-circulating SARS-CoV-2 variants like Delta which necessitates caution as it may lead to increased cases of reinfection.

## Introduction

The emergence of SARS-CoV-2 Variants of Concerns (VOCs) has caused the most severe crisis and challenge to the public health worldwide. The situation has worsened yet again with the recent emergence of highly mutated and super spreading, Omicron variant (B.1.159) in the month of November, 2021 [1]. As of January 6^th^ 2020, the variant has been reported from 149 countries [1]. Increased incidences of COVID-19 cases are being reported from countries where Omicron is becoming dominant variant like the United Kingdom, Denmark and United States of America [1,2]. Millions of people are getting infected with Omicron daily, hence there is an urgent need to deduce the properties of this VOC.

Omicron variant possess around 26 to 32 mutations in the spike protein including mutations associated with increased receptor binding and immune escape property [3]. Three sub lineages have been identified for Omicron variant i.e., BA.1, BA.2 and BA.3, of which BA.1 contributes to the maximum cases reported around globe [4,5]. The preliminary data available on the phenotypic characteristics of the Omicron variant shows increased transmissibility, risk of reinfection, reduced disease severity and need for hospitalization [1]. More data needs to be assessed accounting to the population immunity and other co-morbid conditions to understand the severity of Omicron infection in humans. Apart from the unique mutations identified in the spike protein of this VOC, an additional amino acid change R346K in the variant is being monitored now [6]. In India, R346K has been observed in ∼33% of SARS-CoV-2 Omicron BA.1 variant sequences submitted in GISAID database till 14^th^ January, 2022 [7].

Multiple animal models have been characterized for SARS-CoV-2 and have been used in preclinical studies and other allied research [8]. Syrian hamsters seems to be an appropriate model for SARS-CoV-2 in terms of ACE-2 binding and also the effective replication of the virus in respiratory tract producing pneumonia [9,10]. The model have been effectively used for the studies on the vaccines, antivirals, pathogenesis, transmissibility, virus shedding and disease severity associated with SARS-CoV-2 VOC’s [9,11–21]. Recently, few studies have been published on the pathogenesis aspect of Omicron variant in mice and Syrian hamsters in comparison with other VOC’s demonstrating decreased severity of infection [22–27].

After multiple failed attempts of isolating the Omicron variant in cell lines reported to be susceptible for SARS-CoV-2, we attempted virus isolation using the Syrian hamster model. The virus grew to high titres in the nasal turbinates’ of the model [28]. The deep sequencing of the specimens revealed only 0.0066% difference in the hamster passage sequence with that of the clinical sample with an additional Q19E substitution in the M gene. Apart from the signature mutations of the Omicron variant, the isolate also possessed the R346K mutation. All the available animal studies till date has used the Omicron variant of BA.1 sub lineage without the R346K mutation. Here, we report the characterization of Omicron variant of BA.1 sub lineage with R346K mutation in Syrian hamster model for its pathogenicity, virus shedding pattern and immune response.

## Methods

### Virus

SARS-CoV-2 Omicron variant (GISAID accession no: EPI_ISL_8542938) belonging to the BA.1 sub lineage isolated from nasopharyngeal swab of a COVID-19 patient by inoculating into Syrian hamster was used for the study [28]. The virus titer was found to be 10^5^ TCID_50_/ml on titration in Vero CCL81 cells as per the Reed and Muench method. On deep sequencing following aminoacid changes were found in the isolate. (NSP5_P132H,Spike_T95I,Spike_K417N,Spike_S373P,Spike_Q493R,Spike_N969K,Spike_H655Y,Spike_N856K,N_R203K,Spike_S371L,NSP3_A1892T,Spike_Q954H,Spike_G339D,N_P13L,Spike_N501Y,NSP14_I42V,Spike_P681H,M_Q19E,Spike_N440K,NSP4_T492I,Spike_S375F,Spike_Q498R,Spike_G446S,Spike_N679K,Spike_N764K,Spike_S477N,Spike_Y505H,NSP3_K38R,Spike_R346K,NSP6_I189V,Spike_T547K,M_D3G,Spike_D796Y,Spike_E484A,N_G204R,Spike_T478K,E_T9I,Spike_L981F,M_A63T,NSP12_P323L,Spike_D614G,Spike_ G496S) [28].

### Hamster experiments

The experiment was approved by the Institutional Animal Ethics Committee of ICMR-National Institute of Virology (ICMR-NIV), Pune. The guidelines of the Committee for the Purpose of Control and Supervision of Experiments on Animals (CPCSEA), Government of India were followed during the experiment. Twenty-four Syrian hamsters (either sex) of 5-6-week age procured from a facility of ICMR-NIV, Pune licensed by CPCSEA were used for the study. Individually ventilated cage system was used for animal housing in the maximum containment facility with *ad libitum* food and water. Hundred microlitre of a virus dose of 10^4^ TCID_50_ was used to infect the twenty hamsters intranasally. A group of age matched hamsters (n=4) were kept till 14 days as uninfected control for the study. The hamsters were monitored for 14 days for body weight changes. Throat swab (TS) samples (n=8) with thin nylon flocked swabs, nasal wash (NW) samples (n=8) with sterile phosphate buffered saline (PBS) and freshly voided feces samples (n=8) were collected in 1 ml viral transport media (VTM) on days 1,3,5,7 and 10. On the 14^th^ day, samples were collected from the remaining 4 hamsters. Four hamsters/group were euthanized on days 3, 5, 7, 10- and 14-days post infection (DPI). Organ samples collected during the necropsy were weighed and homogenized using a tissue homogenizer (Qiagen, Germany) and were used for viral load estimation. Lung tissues were examined for their gross changes and fixed in 10 % neutral buffered formalin for histopathological evaluation (Fig.1).

**Fig. 1:**
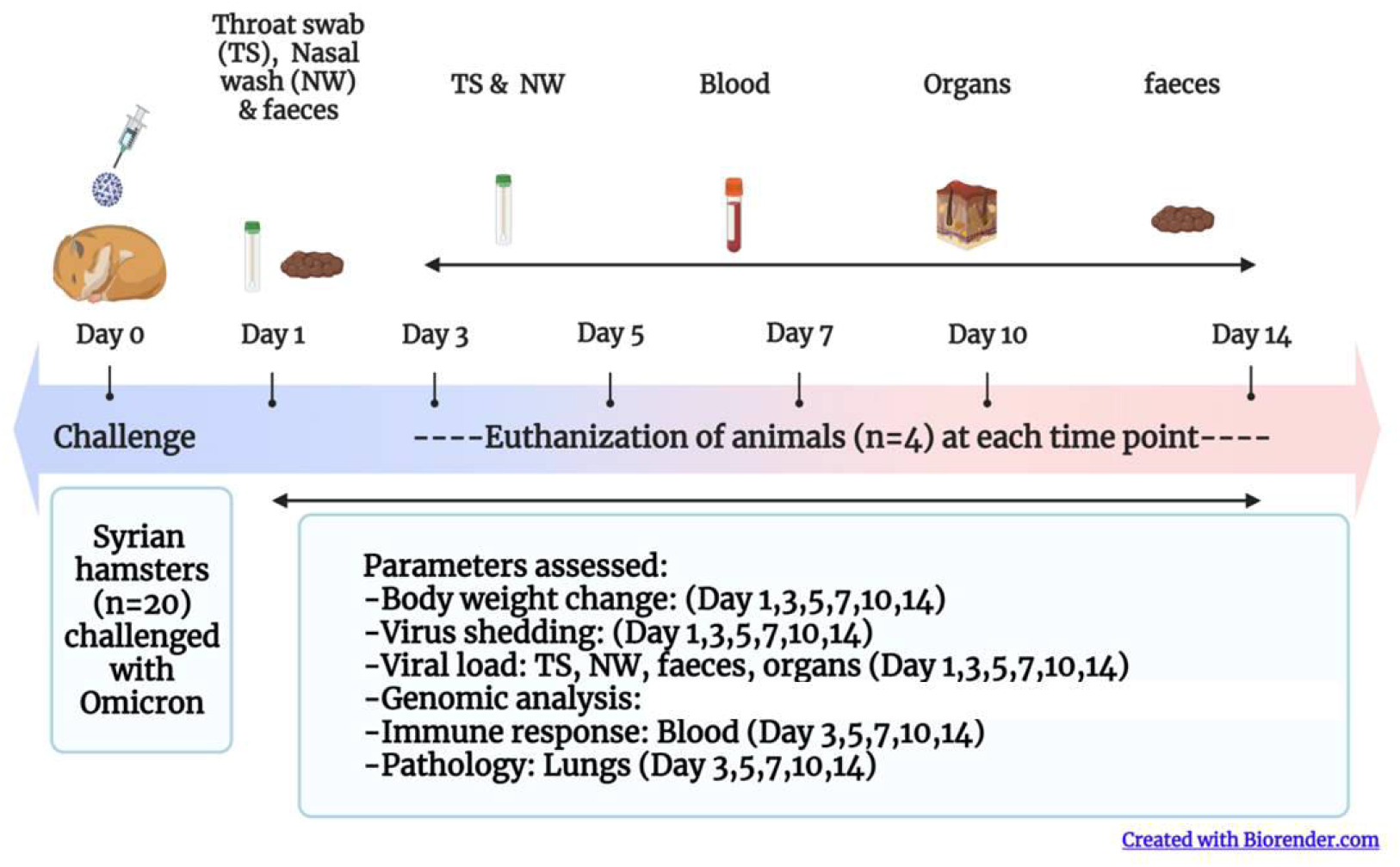
Study design. a) Summary of the study

### Viral load estimation

RNA extraction was performed using the MagMAX™ viral/pathogen nucleic acid isolation kit (ThermoScientific, USA) as per the manufacturer’s instructions. RT-qPCR was performed further using published primers for E gene and N gene for genomic and sub genomic RNA (sgRNA) respectively [29,30]. The lungs, nasal turbinates and NW samples were titrated for the live virus in Vero CCL-81 cells (ATCC, USA) to determine the presence of replication competent virus as described earlier [15]. For titration, 10 fold dilutions of samples were added onto cell monolayer in a 24 well plate and were incubated for one hour. After removal of the media, the cell monolayer was washed with phosphate buffered saline (PBS). The cells were further incubated with maintenance cell culture media containing serum (Gibco, USA) in a CO_2_ incubator. The cells were observed daily for any cytopathic effects (CPE). On observation of CPE, the supernatant from the wells were tested by RT-qPCR for confirmation.

### Anti-SARS-CoV-2 IgG ELISA

The hamster serum samples were tested for antibodies using anti-SARS-CoV-2 hamster IgG ELISA [32] and S1-Receptor Binding Domain (RBD) ELISA. For S1-RBD ELISA, 96-well polystyrene microtitre plates (Nunc, Germany) coated with 1.5 μg/well concentration of S1-RBD protein (Labcare, India) was used. Five percent skimmed milk in 1 X PBS with 0.1 per cent Tween-20 (PBST) (Sigma-Aldrich, USA) was used blocking the wells. The plates were washed thrice after blocking for 2 hours at 37°C. The serum samples were added to the plates and were further incubated for an hour. After washing the plates, 100μl of anti-hamster IgG horseradish peroxidase (HRP) (Thermofisher, USA) was added to each well and incubated for 30 minutes at 37°C. 3, 3’, 5, 5’-tetramethylbenzidine (TMB) was used as substrate and the reaction was terminated using 1 N sulphuric acid. The optical density (OD) was measured using an ELISA reader at 450 nm and the cut off was set at twice the OD value of the negative control.

### Plaque Reduction Neutralization test

The neutralization assay was performed for the serum collected on each time point (3, 5, 7, 10 and 14 DPI) against B.1 variant and Delta, Alpha, Beta, Delta and Omicron VOCs of SARS-CoV-2 as described earlier [31].

### Cytokine response

Cytokine ELISA was performed using commercially available kit (Immunotag, USA. The levels of IFN-γ, IL-4, IL-6, IL-10 and TNF-α were estimated. The cytokine levels were compared with that of serum cytokine levels of 3 healthy control animals.

### Histopathology and immunohistochemistry

Lungs samples fixed in formalin were processed for histopathology by hematoxylin and eosin staining [33]. Coded tissue samples were blindly scored by a pathologist. Each section was scored for vascular changes, bronchiolar changes, inflammatory cell infiltration, emphysema, oedema, hyaline changes and alveolar pneumocyte changes [17]. Each lesion was given a score of 0 to 1 and the cumulative score was plotted.

For immunohistochemical evaluation, duplicate sections were used as that of histopathology. Lung tissue sections were rehydrated and 0.3% hydrogen peroxide in methanol was used for antigen retrieval. Polyclonal anti-SARS-CoV-2 mouse serum (generated in the laboratory using inactivated SARS-CoV-2 antigen in mice) was used as the primary antibody. The sections were incubated with primary antibody (1: 400 dilution) for 60 minutes and washed with PBS. Goat anti-mouse horse radish peroxidase antibody (Dako, USA) and 3, 3’-diaminobenzidine (DAB) tetrahydrochloride substrate were used as secondary antibody and for detection respectively.

### Next Generation Sequencing

Viral RNA was extracted with QIAmp Viral RNA extraction kit (Qiagen, Germany). Qubit® 2.0 Fluorometer (Invitrogen, USA) was used for quantifying the RNA with the Qubit RNA High Sensitivity (HS) kit (Thermofisher, USA). The RNA library was prepared using the TruSeq Stranded mRNA LT Library preparation kit (Illumina, USA). The library was quantified, normalized (1.8 picomolar) and loaded on to the Illumina Miniseq Next Generation Sequencing platform. The reads generated were mapped along with the SARS-CoV-2 Wuhan isolate (Accession No.: NC_045512.2) using the reference-based assembly method of the CLC Genomics Workbench v.20. The nucleotide variations were identified using the basic variant tool as implemented in the CLC Genomics Workbench v.20.

### Data analysis

The experimental data was analyzed using Graph pad Prism version 9.2.0 software. Non parametric Mann Whitney tests were used for the body weight comparison of the infected with the control animals and Kruskal Wallis tests was used for the comparison of neutralization data. A p-value less than 0.05 were considered as statistically significant.

## Results

### Omicron infection reduced body weight gain in hamsters

The body weight gain in hamsters infected with Omicron variant was less compared to the control animals (Fig. 2A). Percent mean body weight difference of -3.2% (*p*=0.004 vs control), -0.53% (*p*=0.016 vs control) and - 0.08% (*p*=0.004 vs control) were observed on day 3, 5 and 7 DPI in Omicron infected animals whereas control group showed an increase of 4.17%, 9.8% and 15.25 % on day 3, 5 and 7 DPI respectively. Thereafter an increase in the mean body weight was seen in the Omicron infected hamsters which was again significantly lesser in comparison to the animals from the uninfected control group.

**Fig. 2:**
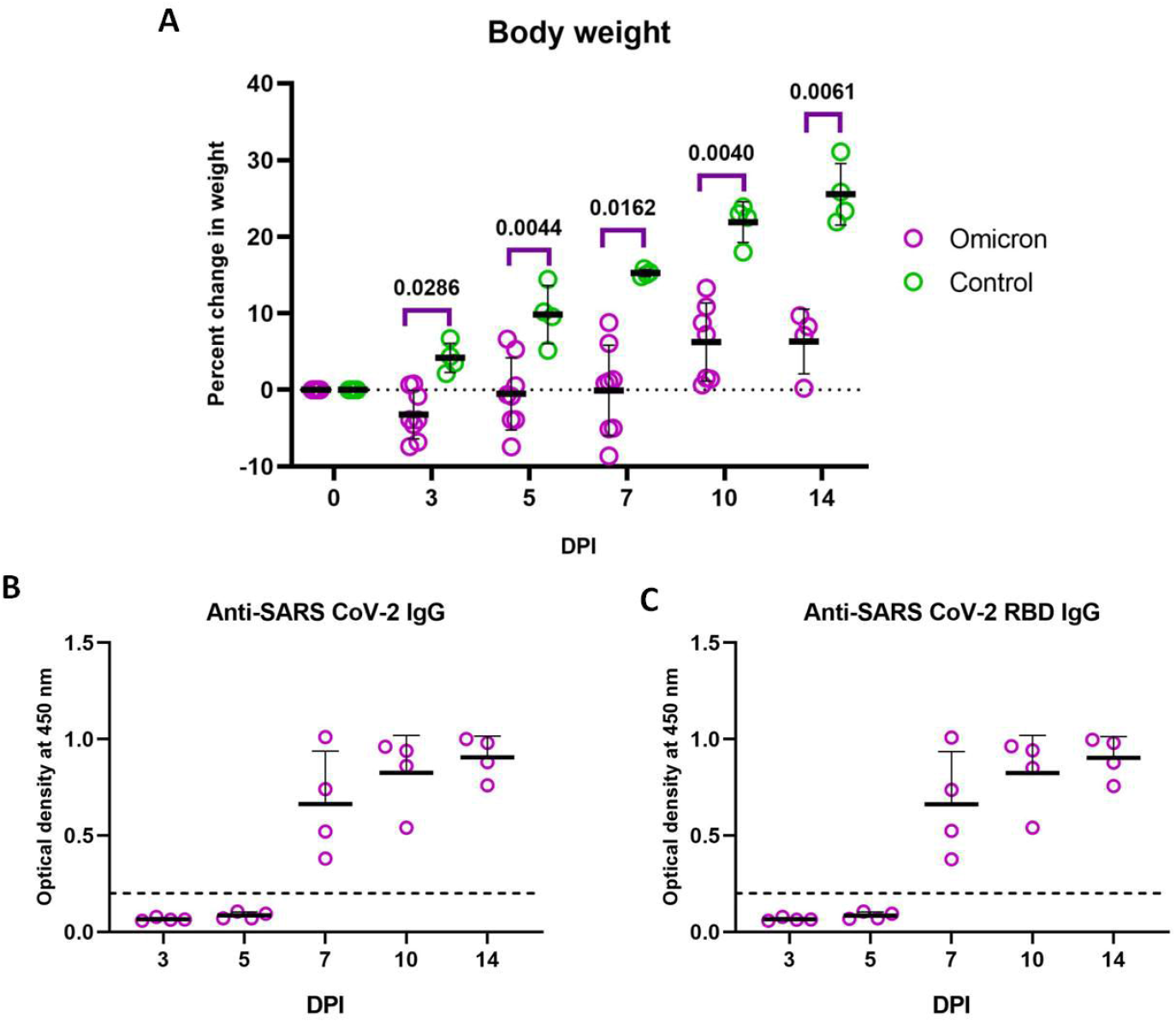
Percent body weight change and IgG response in hamsters post infection. **A)** Scatter plot showing the body weight changes in hamsters after Omicron infection in comparison to that of age matched control animals. [*Mann Whitney test*, Omicron (n=8 on 3, 5, 7 and 10 DPI and n=4 on 14 DPI) *vs* uninfected control (n=4)]. **B)** Anti-SARS-CoV-2 inactivated antigen IgG response in hamsters post infection by ELISA. **C)** Anti-SARS-CoV-2 RBD antibodies in hamsters post infection by ELISA.

### Immune response in hamsters after infection

The IgG antibodies could be detected from 7 DPI in hamsters (Fig. 2B,C). The neutralizing antibodies (NAbs) against Omicron variant was detected in infected hamsters from 5 DPI with geometric mean titre (GMT) of 98.30, 2170.40, 932.62 and 953 on 5,7,10 and 14 DPI respectively (Fig. 3A). The neutralizing ability of the antibodies against other variants was found to be substantially reduced. Against Alpha variant, GMT was 136.6 on 14 DPI and no neutralization was observed against the B.1, Beta and Delta variants (except one hamster sera) (Fig. 3B). No significant elevation was seen in the IFN-γ, IL-4, IL-10 and TNF-α levels in sera of infected animals compared to the control animals. The IL-6 and IFN-γ levels showed mild increase on 5 DPI in the Omicron infected animals although it was not statistically significant (Supplementary Figure 1).

**Fig. 3:**
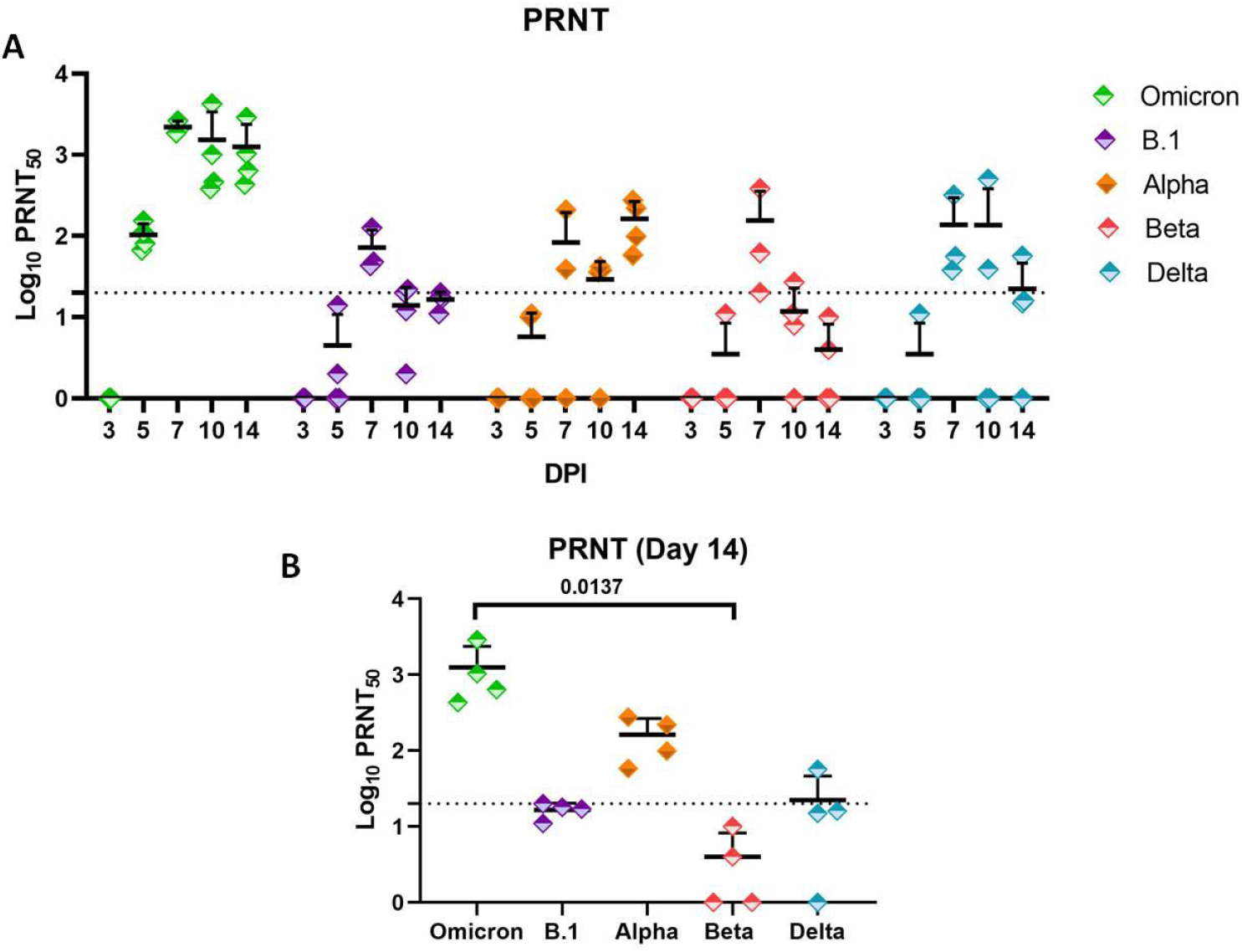
Neutralizing antibody levels in hamsters post Omicron infection. **A)** Scatter plot showing neutralizing antibody levels in Omicron infected hamsters on 3,5,7,10 and 14 DPI against Omicron, B.1, Alpha, Beta and Delta variants. **B)** Scatter plot showing neutralizing antibody levels on day 14 against Omicron, B.1, Alpha, Beta and Delta variants. [*Kruskal Wallis test*, p=0.0137, Omicron vs Beta].

### Virus shedding following Omicron infection in Syrian hamsters

Viral gRNA was detected in the NW, TS and faeces samples of Omicron infected hamsters and the period of detection varied among samples (Fig. 4A-C). The NW and TS samples showed presence of viral gRNA till 14 DPI in all hamsters and till 5 DPI in faeces samples. The highest mean gRNA load [mean ± standard deviation (SD) = 4.6 × 10^8^ ± 3.6 × 10^8^ genome copies/ ml] was observed on 3 DPI in the NW samples, which showed decrease in the genome copies on further days. The TS samples showed highest mean genome copies (mean ± SD= 2.1 × 10^8^ ± 3.6 × 10^8^) on 1 DPI and decreased further. Subgenomic RNA detection window was further lower for all samples ie., till 10, 7 and 5 DPI in NW,TS and faeces samples respectively (Fig. 4D-F). The sgRNA levels in the samples also showed a similar trend as that of gRNA. The mean live virus load in the NW sample was found to be 1.86 × 10^3^ TCID50/ml on 1 DPI, 2.08 × 10^2^ TCID50/ml on 3 DPI and 1.54 × 10^2^ TCID50/ml on 5 DPI (Fig. 4G).

**Fig. 4:**
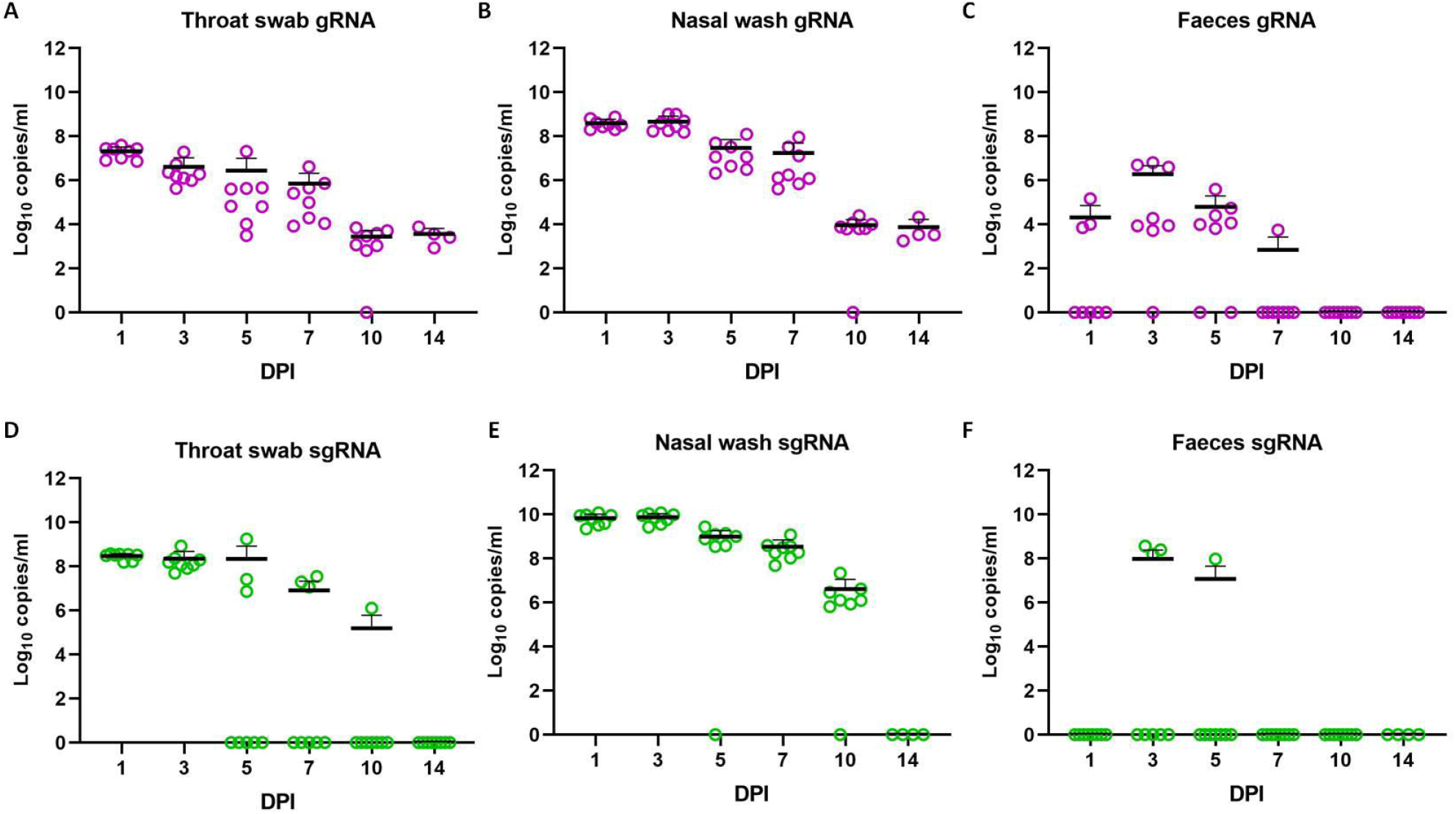
Viral shedding in hamsters after Omicron infection. **a)** Scatter plot showing viral gRNA load in A) Throat swab B) Nasal wash C) Faeces and viral sgRNA load in D) Throat swab E) Nasal wash F) Faeces in hamsters on 1,3,5,7,10 and 14 DPI post Omicron infection. G) Viral load in the nasal wash samples of hamsters.

### Viral load in the organs

Among the organs tested for viral gRNA and sgRNA, respiratory organs (nasal turbinates, trachea, bronchi and lungs) showed consistent detection on all the sampled days. All the other organs (brain, liver, heart, spleen intestine and kidney) were negative except that of one hamster on 3 DPI which showed viral gRNA and sgRNA detection in liver, small intestine and serum samples (Figure 5A-H). The highest mean viral RNA load was found on 3 DPI in the respiratory organs. The viral RNA levels showed reduction on further days. Unlike 2 log reduction in viral RNA in nasal turbinates, lungs showed lesser decrease on 5 DPI. The mean viral gRNA levels on 3 DPI was 1.09 × 10^10^ ± 1.36 × 10^10^ (mean ± SD) and on 5 DPI was 2.12 × 10^9^ ± 3.9 × 10^9^ (mean ± SD) copies/ml in lung samples. On virus titration a mean titre of 1.9 × 10^4^ TCID50/ml on 3 DPI and 5.49 × 10^2^ TCID50/ml on 5 DPI in the lungs sample and 3.71 × 10^3^ TCID50/ml on 3 DPI in nasal turbinate were observed. No viral titre could be detected in the lungs and nasal turbinate samples on further days (Fig. 5I).

**Fig. 5:**
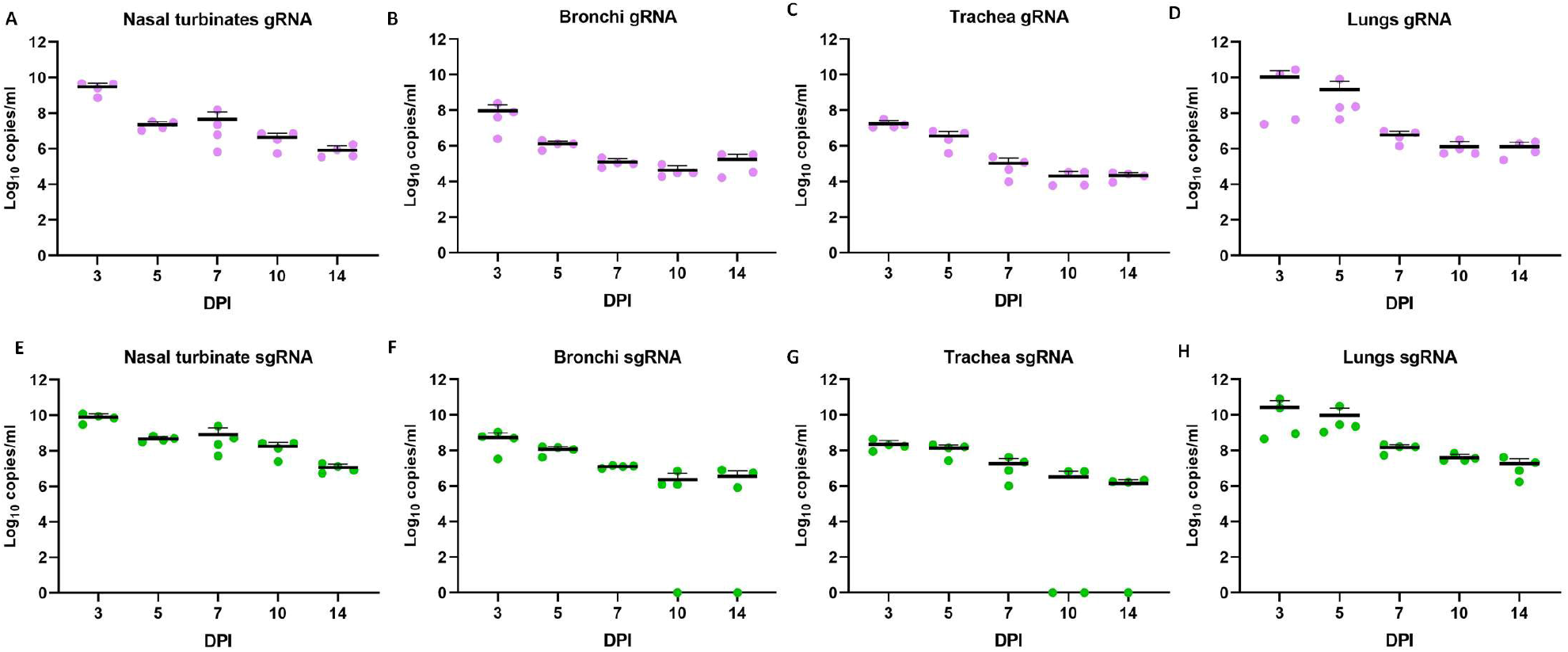
Viral RNA load in organs of hamsters after Omicron infection. Scatter plot showing viral gRNA load in A) Nasal turbinates B) trachea C) Bronchi and D) Lungs. Viral sgRNA load in E) Nasal turbinates F) Trachea G) Bronchi and H) Lungs in hamsters post Omicron infection. Viral load (TCID_50_) in I) Nasal turbinates and J) Lung samples of hamsters.

### Lung pathological changes in hamsters after infection

Gross changes like congestion and haemorrhages were visible in the lungs of hamsters from 3 DPI which became pronounced on 5 and 7 DPI (Figure 6A-H). Histopathologically early changes observed were congestion and bronchial epithelial necrosis with exudates on 3 DPI which progressed to diffuse alveolar damage on 5 and 7 DPI. Emphysema, haemorrhages, hyperplasia of alveolar epithelial cells, oedema and mononuclear cell infiltration were also observed in the alveolar parenchyma. Peribronchial and perivascular inflammatory cell infiltration was found during 7 DPI. By 14 DPI, except in case of a single hamster, the lesions observed were found to be resolved (Fig. 6I-L). Viral antigen could be detected in the bronchial epithelium, alveolar epithelial cells and macrophages. The immunostaining was prominent in the bronchiolar epithelial cells on 3 DPI and in both bronchiolar and alveolar on 5 and 7 DPI. On further days only faint or focal immunostaining was observed in the alveolar epithelial cells (Fig. 6M-O).

**Fig. 6:**
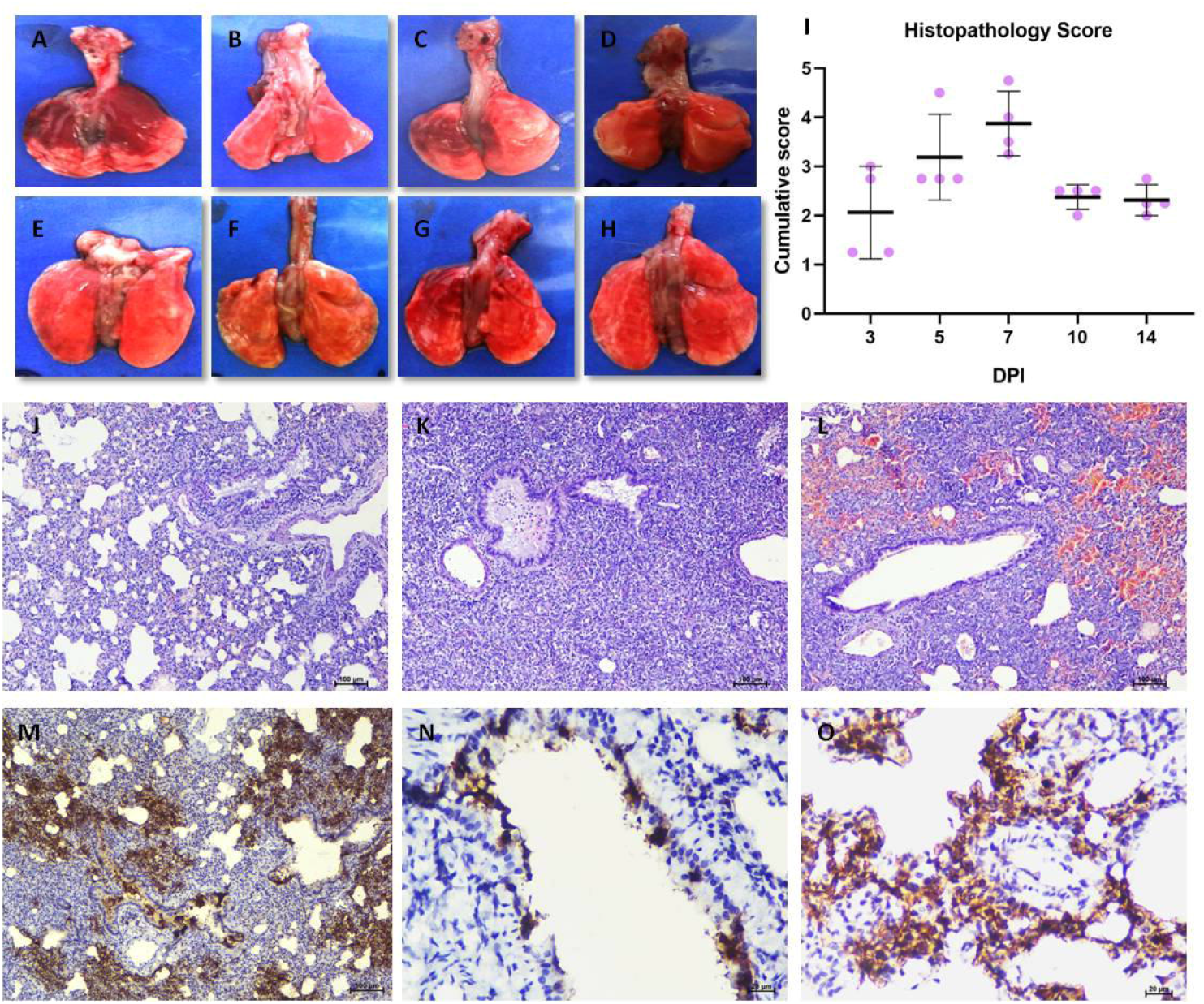
Pathological changes in hamsters after omicron infection. Lungs of hamsters on 5 DPI (A-D) and 7 DPI (E-H) showing varying degree of consolidation and haemorrhages. I) Cumulative histopathological score of lung lesions observed on 3, 5, 7, 10 and 14 DPI in hamsters, which was scored out of 8. J) Lung sections showing exudates in bronchiole with peribronchiolar cellular infiltration and alveolar consolidation, H& E. K) Lungs sections showing bronchioles filled with exudates and denuded epithelial cells as well as inflammatory cells, diffuse consolidation and alveolar septal thickening in the parenchyma as well as peribronchial and perivascular mononuclear cell infiltration. L) Lung section showing diffuse alveolar haemorrhages, peribronchial and perivascular mononuclear cell infiltration and bronchiolitis. M) Bronchiolar and alveolar epithelial cell showing intense and diffuse immunostaining on 5 DPI by immunohistochemistry, DAB. N) Bronchiolar epithelial cells showing intense immunostaining on 5 DPI by immunohistochemistry, DAB O) Alveolar epithelial cells showing intense immunostaining on 5 DPI by immunohistochemistry, DAB.

### SARS-CoV-2 Variant analysis

We performed the variant analysis of the Omicron virus isolate, inoculum as well as the Omicron infected lung samples collected on 3, 5, 7 and 10 DPI. The analysis showed no additional mutations in the genome other than that of the virus isolate (Fig. 7).

**Fig 7:**
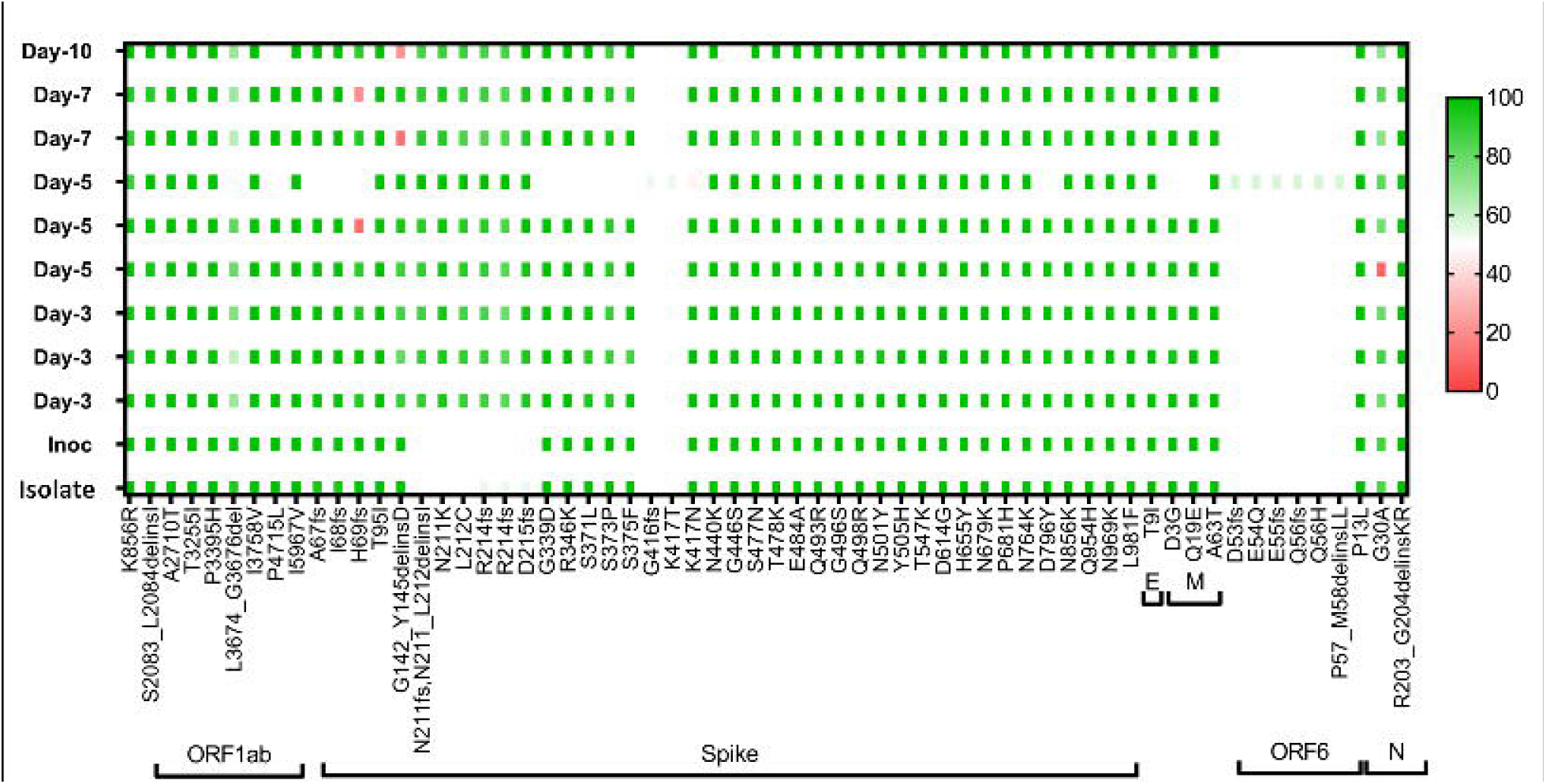
SARS-CoV-2 variant analysis. The plot showing the amino acid substitutions in the virus inoculum and lung samples post infection on 3, 5, 7 and 10 DPI.

## Discussion

The Omicron variant is spreading worldwide at an alarming rate and the researchers across the globe are putting their best efforts to understand the characteristics of the virus. The Omicron variant we used in this study had an R346K substitution unlike other studies available in public domain. Though specific role of this mutation is not well studied but the substitution is being monitored.

The disease characteristics of the Omicron variant observed here are similar to those reported for other SARS-CoV-2 variants like Alpha, Beta, Kappa and Delta in hamsters by earlier studies except that of the body weight loss [9,19,32,33]. Our finding corroborated with the few recent studies which also reported the limited or negligible body weight loss in hamsters post infection with Omicron variants [22–26]. The age and virus dose can affect the body weight loss and disease severity in hamsters post SARS-CoV-2 infection [20, 32]. We have used 5-6 week old hamsters and other published studies on Omicron infection in Syrian hamsters have also used animals of the age range of 4 to 8 weeks except Ryan et al., have used 20-22 week animals [22]. Even high Omicron virus doses could not result in body weight loss in hamsters [23]. In the present study we have used only a single dose for virus infection.

The viral RNA shedding and live virus clearance observed here was similar to earlier VOCs studied in hamster model [21,34]. Replication competent virus could be detected in NW till 5 DPI. The clearance of virus was observed by 1 week in hamsters. Virus shedding period found in human Omicron cases are also suggesting that infectious period is lesser than 10 days since symptom onset [35].

We observed moderate to severe changes in the lungs of hamsters infected with Omicron. In contrary to our findings, a recent *ex vivo* study reported that Omicron infectivity of lungs is lower in comparison to the Delta variant [36]. Lung pathology observed in the Omicron studies in hamsters also reports that lung disease is milder in comparison to the ancestral strains [23,25,26]. The virus dose used in each study differed and the time point used for assessing histopathological disease score also varied. Here we have selected multiple time points to understand the difference in lung disease severity. But we have used only a single virus dose for infection and we did not assess the comparative severity with other variants. In comparison to our earlier studies of SARS-CoV-2 variants in Syrian hamster model, the lung lesions observed with Omicron was prominent. We also observed higher viral load in the lungs similar to that of upper respiratory tract in hamsters. The virus titration showed early virus clearance from the nasal turbinates by day 5 and from lungs by day 7. We found no evidence that Omicron produce less severe lung disease or more predilection to upper respiratory tract in comparison to lung as reported earlier [20, 22, 23, 24].

Neutralizing antibodies could be detected from 5^th^ DPI in Omicron infected hamsters, which showed an increase in titre on further days similar to other VOCs [19]. The serum samples of hamster infected with Omicron showed poor neutralization against other VOCs tested like Alpha, Beta and Delta variants. As Omicron variant harbours around 26-32 mutations in the spike protein RBD and NTD, which are the most antigenic/ immunodominant region of the virus, immune evasion was speculated for the variant [37]. It harbours N501Y, K417N, T478K, E484A which are amino acid substitutions linked with immune escape[38]. In addition virus isolate used in study also possess R346K mutation which is also present in the Mu variant known for immune escape [39]. So far, neutralization studies on convalescent sera of Omicron infected patients and animal models are not available. The studies published on the neutralization potential of the convalescent sera samples of COVID-19 recovered as well as vaccinated individuals showed poor neutralization against the Omicron variant [40,41]. Our preliminary data suggests that humoral immune response generated by Omicron infection may not confer much protection to other current circulating variants like Delta. The other studied variants like B.1, Beta and Alpha variants have almost disappeared from circulation globally. This suggests constant monitoring of immunized population for their antibody levels and studies on vaccine efficacy against emerging variants.

Here, we have characterized an Omicron isolate of BA.1 sub lineage with R346K mutation in Syrian hamster model. We observed limited body weight gain, viral replication in the upper and lower respiratory tract and interstitial pneumonia in Syrian hamsters following infection with the Omicron variant. Moreover, the NAbs generated after Omicron infection showed substantial reduction or negligible neutralization against other variants like B.1, Alpha, Beta and Delta. The presented data provides new insight into the neutralization potential and cross protection of Omicron variant against other VOCs.

## Contributors

PD Yadav conceived and designed the study. S Mohandas performed the animal experiments. A Shete co-ordinated and performed the real time PCR assays, sample processing and testing. G Sapkal and G Deshpande performed the PRNT assays. D Nyayanit performed the statistical analysis. R Jain and A Shete performed the ELISA based experiments. M Kadam and A Kumar performed the daily sampling and sample processing. C Mote performed the histopathological and immunohistochemical processing and examination. P D Yadav, and S Mohandas analysed and interpreted the results. S Mohandas and DY Patil wrote the preliminary manuscript draft. All the authors substantially revised the manuscript.

## Role of funding source

The study sponsor has no role in study design, analysis, interpretation of data, in the writing of the report and in the decision to submit the paper for publication.

## Declaration of Interests

The authors declare no competing financial interests.

## Acknowledgments

This study was supported by Indian Council of Medical Research as an intramural grant (COVID-19) to ICMR-National Institute of Virology, Pune. Authors acknowledge the support of Prof. Priya Abraham, Director, ICMR-NIV, Pune. The authors also acknowledge the support received from the laboratory team of Maximum Containment Facility of ICMR-NIV, Pune ie., Mr Prasad Sarkale, Mr Hitesh Dighe and Mr Rajen Lakra for tissue culture support, Mr Annasaheb Suryawanshi and Mr Kundan Wakchaure for laboratory animal care and sample processing, Mrs Kaumudi Kalele, Ms Jyoti Yemul, Ms Pranita Gawande, Ms Poonam Bhodke, Mrs Priyanka Waghmare for sample testing and Mrs Triparna Majumdar, Mrs Savita Patil, Mr Yash Joshi and Ms Manisha Dudhmal for support in sequencing and analysis.

## Data Sharing Statement

All the data pertaining to the study are available in the manuscript or in the supplementary materials.

## Figure legends

**Supplementary figure 1. Serum cytokine levels in hamsters post Omicron infection**. A) IFN-Gamma B) IL-4 C) IL-6 D) IL-10 E) TNF-alpha

## Notes

### Competing Interest Statement

The authors have declared no competing interest.

