## Supplementary figures and images for "Pathogenicity of SARS-CoV-2 Omicron in Syrian hamsters and its neutralization with different Variants of Concern"

### Supplementary figure 1

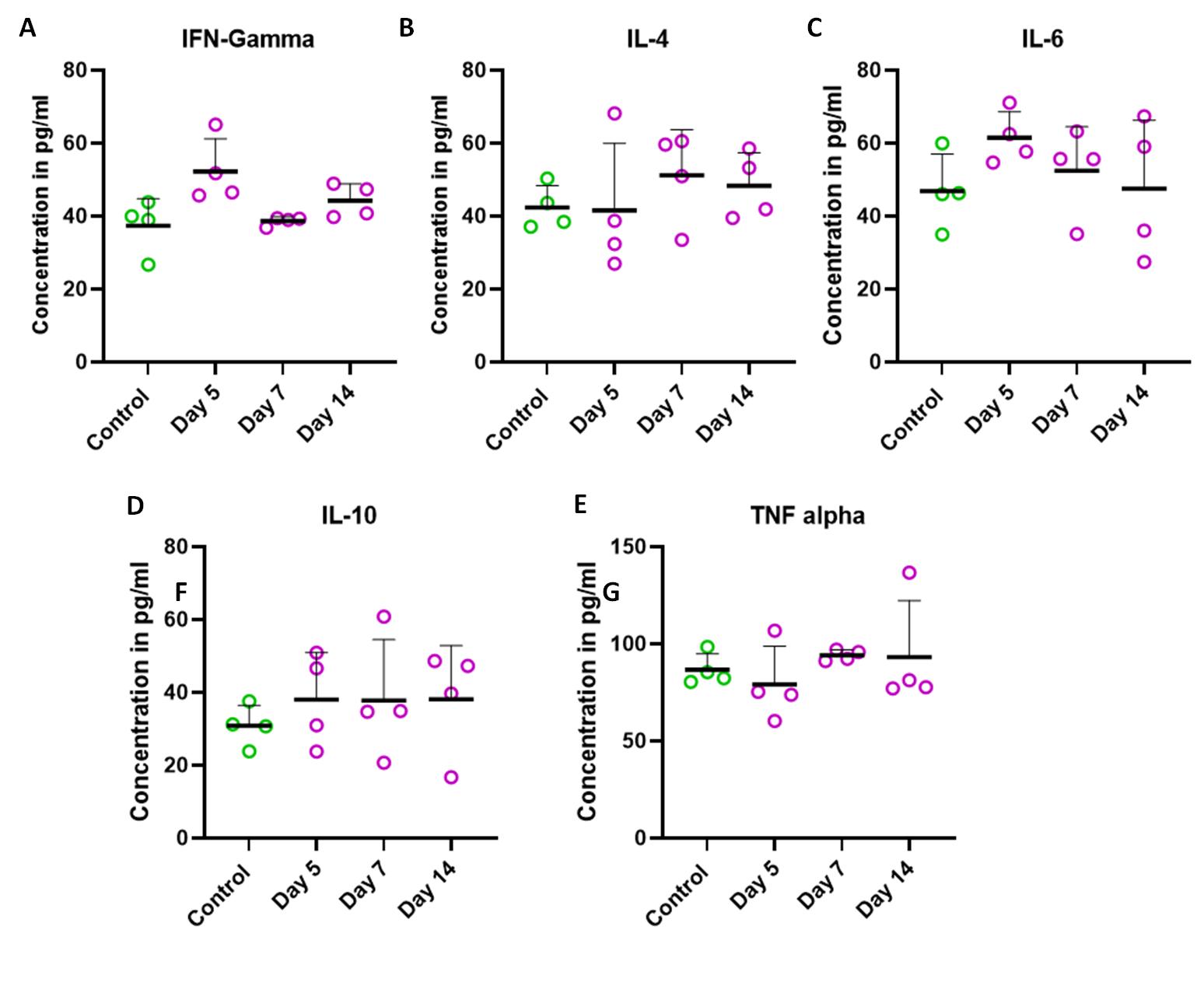
